# Fish diversity in a human-impacted coastal lagoon in Central Peru (~10°78’ °S)

**DOI:** 10.1101/2025.11.07.687100

**Authors:** Víctor Aramayo

## Abstract

Eight fish species were recorded in a heavily impacted coastal lagoon on the central Peruvian coast. Despite anthropogenic pressures such as waste disposal and habitat fragmentation, the lagoon still supports a typical estuarine fish assemblage, reflecting the resilience of this ecosystem. A total of 2357 individuals representing eight species, seven families, and five orders were collected. The identified species were *Bryconamericus peruanus, Dormitator latifrons, Lebiasina bimaculata, Poecilia reticulata, Ctenogobius sagittula, Mugil cephalus, Cichlasoma nigrofasciatum*, and *Aequidens rivulatus*. Among these, *B. peruanus* and *D. latifrons* were dominant, comprising more than half of all captures. Most individuals were juveniles, confirming that the Totoral Lagoon functions as a nursery ground that provides favorable conditions for growth and recruitment, including abundant food, vegetation cover, and stable physicochemical parameters. Fish were concentrated in shallow, vegetated, muddy zones where freshwater and marine influences converge. These areas serve as refugia and feeding grounds, particularly for species that complete early life stages within the lagoon. Native taxa such as L. bimaculata, *D. latifrons*, and *C. sagittula* occurred in lower densities, mostly in low-salinity areas with fine sediments, indicating the persistence of localized freshwater microhabitats. Overall, the structure and composition of the ichthyofauna reveal a marked ecological gradient from freshwater to brackish zones, reinforcing the lagoon’s function as a key transitional ecosystem along the Peruvian coast.

## Introduction

Freshwater ecosystems, such as coastal lagoons, wetlands, and mangroves that connect to the sea, are key areas for the reproduction and growth of many fish species and other organisms that utilize the unique features of these environments [1–3]. These aquatic grounds vary considerably in their connections with the sea, and have multiple trophic interactions with strong relationships between nekton and the surface sediment-dwelling organisms [4,5], all of which make these fragile ecosystems in coastal areas highly dynamic [6]. In some cases, it has been suggested that these ecological transition ecosystems (ecotones) are key areas for the maintenance of many commercially important coastal species [7]. Fish are among the most representative fauna of these environments, and they are considered as indicators of environmental quality [8]. Their community structure and diversity might reflect the gradual ecosystem recovery (or resilience) in cases of human-induced pollution or natural disturbance changes in the habitat [9].

Coastal lagoons represent dynamic ecotones where marine and continental processes converge, creating highly productive and biologically complex environments. These semi-enclosed systems act as critical nursery, feeding, and refuge habitats for numerous fish species, mediating energy and nutrient fluxes between terrestrial catchments and adjacent coastal waters [10]. Their ecological functionality depends strongly on the degree of connectivity with the sea, which regulates salinity gradients, sediment transport, and benthic–pelagic coupling processes that sustain local productivity [11,12]. Within such systems, benthic communities, ranging from meiofauna to macroinvertebrates, play pivotal roles in nutrient regeneration, sediment bioturbation, and as prey resources for demersal and juvenile fishes [13].

Despite their ecological significance, coastal lagoons across the southeastern Pacific, particularly along the Peruvian coast, have received limited scientific attention. Human pressures such as artisanal fishing, urban encroachment, eutrophication, and hydrological alteration are increasingly modifying their ecological structure and functioning. Also, selective fishing practices, targeting size classes or specific trophic guilds, can shift community composition and trophic networks, ultimately affecting ecosystem resilience and fishery yields [14,15,5]. Furthermore, episodic climatic perturbations linked to El NiÑo–Southern Oscillation (ENSO) events periodically alter temperature, salinity, and oxygen regimes, reshaping fish assemblages and their benthic prey bases [16,17].

Understanding fish diversity and its ecological determinants in these transitional environments is essential to interpret the responses of coastal ecosystems under intensifying anthropogenic and climatic stress. In this context, the present study provides the first assessment of fish assemblages in a human-impacted coastal lagoon of central Peru (∼11.5°S), examining diversity, ecological affinities, and potential linkages with benthic structure and trophic dynamics. Such baseline information is critical for evaluating habitat integrity, guiding sustainable fishery management, and framing conservation strategies within the broader Humboldt Current System.

## 2. Materials and Methods

Fish sampling was conducted in the Totoral Lagoon (10°78’59” S, 77°75’47” W), a small coastal waterbody located on the central Peruvian coast, north of Lima (Figure 1). The lagoon covers approximately 25.1 hectares and lies about 2.5 km inland from the shoreline. Its width varies between 4.5 and 32 m, and its depth ranges from 1.0 to 2.8 m. Tidal influence is most evident at night, with the water level increasing by approximately 1 m, particularly during the summer months. The lagoon exhibits a sedimentary gradient extending from coarse boulders in the upper, more continental zone, where a permanent freshwater input occurs, to alternating sandy and muddy sediments in the middle sector, similar to central Peruvian areas [18], and finally to predominantly sandy substrates near the marine outlet. Water movement is essentially unidirectional, creating several lentic microhabitats along its course.

**Figure 1.**
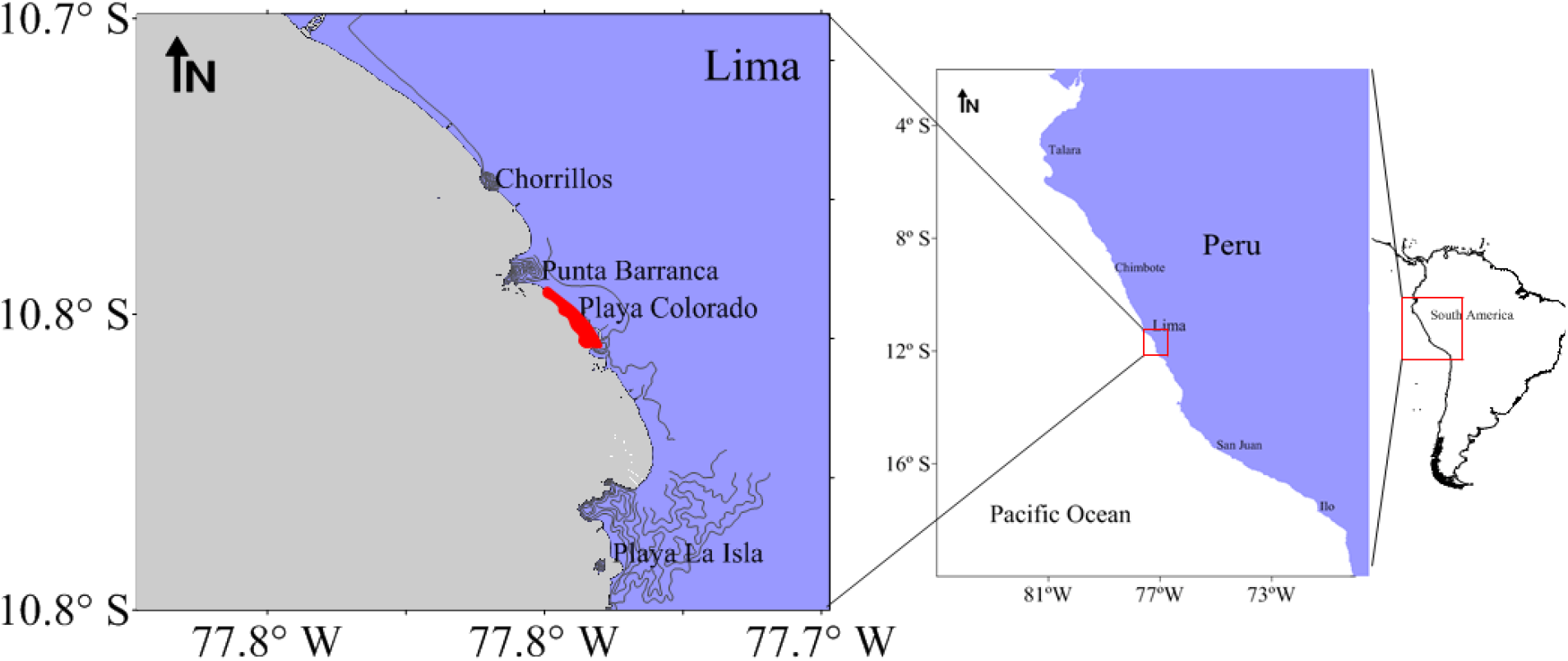
Sampling site, the red shaded area indicates the approximate size and location of the Totoral Lagoon, Central Peru.

Fish specimens were collected from six predetermined sampling stations between July 2003 and March 2004 using shore seine hauls (shore dragging) as the primary collection method. This technique allowed the capture of both benthic and pelagic species inhabiting shallow areas along the lagoon’s margins. Each haul was performed manually, ensuring standardized effort across stations and minimizing habitat disturbance. All collected specimens were preserved and identified to the lowest possible taxonomic level following the diagnostic criteria of Nelson (1994). Field observations and photographic records were used to support taxonomic verification and species quantification. Occasional visual records of *Cichlasoma nigrofasciatum* and *Aequidens rivulatus* were documented during fieldwork but not captured; nonetheless, these species are included in the species inventory presented herein.

## 3. Results and Discussion

The Totoral Lagoon encompasses approximately 25.1 hectares and lies about 2.5 km inland from the coastline. Its width ranges from 4.5 to 32 m, and depth varies between 1.0 and 2.8 m. Water level fluctuations of up to 1 m were observed during nocturnal high tides, particularly in the summer season. The lagoon exhibits a clear spatial gradient in sediment composition: coarse boulders dominate the upper freshwater sector, transitioning into mixed sandy–muddy substrates toward the middle zone, and finally into fine sand near the marine outlet. Water flow follows a predominantly unidirectional pattern, forming localized lentic areas along its course.

A total of fish specimens were obtained from six fixed sampling stations between July 2003 and March 2004 through shore-seine collections. Species were identified to the lowest possible taxonomic level following Nelson (1994). Field observations and photographic records supplemented the identifications and supported the assessment of species occurrence and abundance. Two additional species, *Cichlasoma nigrofasciatum* and *Aequidens rivulatus*, were recorded visually during fieldwork but not captured in the samples; nevertheless, both are included in the species inventory.

A total of 2357 fish specimens were collected during the study period, representing eight species distributed among seven families and five orders (Table 1). The species identified were *Bryconamericus peruanus, Dormitator latifrons, Lebiasina bimaculata, Poecilia reticulata, Ctenogobius sagittula, Mugil cephalus, Cichlasoma nigrofasciatum*, and *Aequidens rivulatus*. Among the species recorded, *Bryconamericus peruanus* and *Dormitator latifrons* were the most abundant, together accounting for more than half of the total individuals captured. These species were present throughout all sampling stations, indicating a broad ecological tolerance to the lagoon’s salinity gradient. In contrast, *Lebiasina bimaculata* and *Poecilia reticulata* were mainly restricted to the inner, freshwater sector of the lagoon, where lower salinity and finer sediments prevailed. The occurrence of *Mugil cephalus* and *Ctenogobius sagittula* was limited to stations near the marine outlet, suggesting an affinity for brackish or coastal conditions.

**Table 1.** Taxonomic list of fishes collected in Totoral Lagoon, Central Peru (10°78’ S).

| Order | Family | Species |
| --- | --- | --- |
| Characiform | Characidae | *Briconamericus peruanus* |
|  | Lebiasinidae | *Lebiasina bimaculata* |
| Perciform | Mugilidae | *Mugil cephalus* |
|  | Eleotridae | *Dormitator latifrons* |
|  |  | *Ctenogobius sagittula* |
|  | Cichlidae | *Cichlasoma nigrofasciatum* |
|  |  | *Aequidens rivulatus* |
| Cyprinodontiform | Poeciliidae | *Poecilia reticulata* |

Although *Cichlasoma nigrofasciatum* and *Aequidens rivulatus* were observed *in situ* during fieldwork, no individuals were captured with the sampling gear. Their presence, however, was confirmed by visual records and photographs, supporting their inclusion in the species inventory. The detection of these cichlids, both typically associated with freshwater or slightly brackish environments, suggests episodic migration events or transient populations influenced by seasonal hydrological connectivity with adjacent water bodies.

Overall, the composition of the ichthyofauna in Totoral Lagoon reflects a mixed assemblage of freshwater, estuarine, and marine-derived species. This pattern indicates the ecological importance of the lagoon as a transitional habitat that supports species exchange between continental and coastal environments.

Most of the specimens collected were juveniles, a pattern commonly observed in coastal lagoons and estuarine systems influenced by tidal dynamics and the continuous inflow of seawater. Juvenile dominance indicates that the Totoral Lagoon functions as a nursery ground, providing favorable conditions for early life stages of multiple species. Such environments offer abundant food resources, shelter from predators, and relatively stable physicochemical conditions that enhance recruitment success. Individuals were mainly concentrated in shallow zones characterized by dense aquatic vegetation and muddy substrates, which are typical features of transitional habitats where freshwater and marine influences converge. These vegetated areas play a key ecological role by serving as refugia and feeding grounds, particularly for species that spawn or complete part of their growth cycle within the lagoon.

Among the native taxa of the central and southern Peruvian coast, *Bryconamericus peruanus* (Figure 2A) was the most prominent species, showing both high abundance and wide spatial distribution within the lagoon. Other native species, such as *Lebiasina bimaculata, Dormitator latifrons*, and *Ctenogobius sagittula* were recorded in lower numbers, generally associated with areas of reduced salinity and finer sediment texture. These species are considered indicators of relatively stable freshwater conditions within the system.

**Figure 2.**
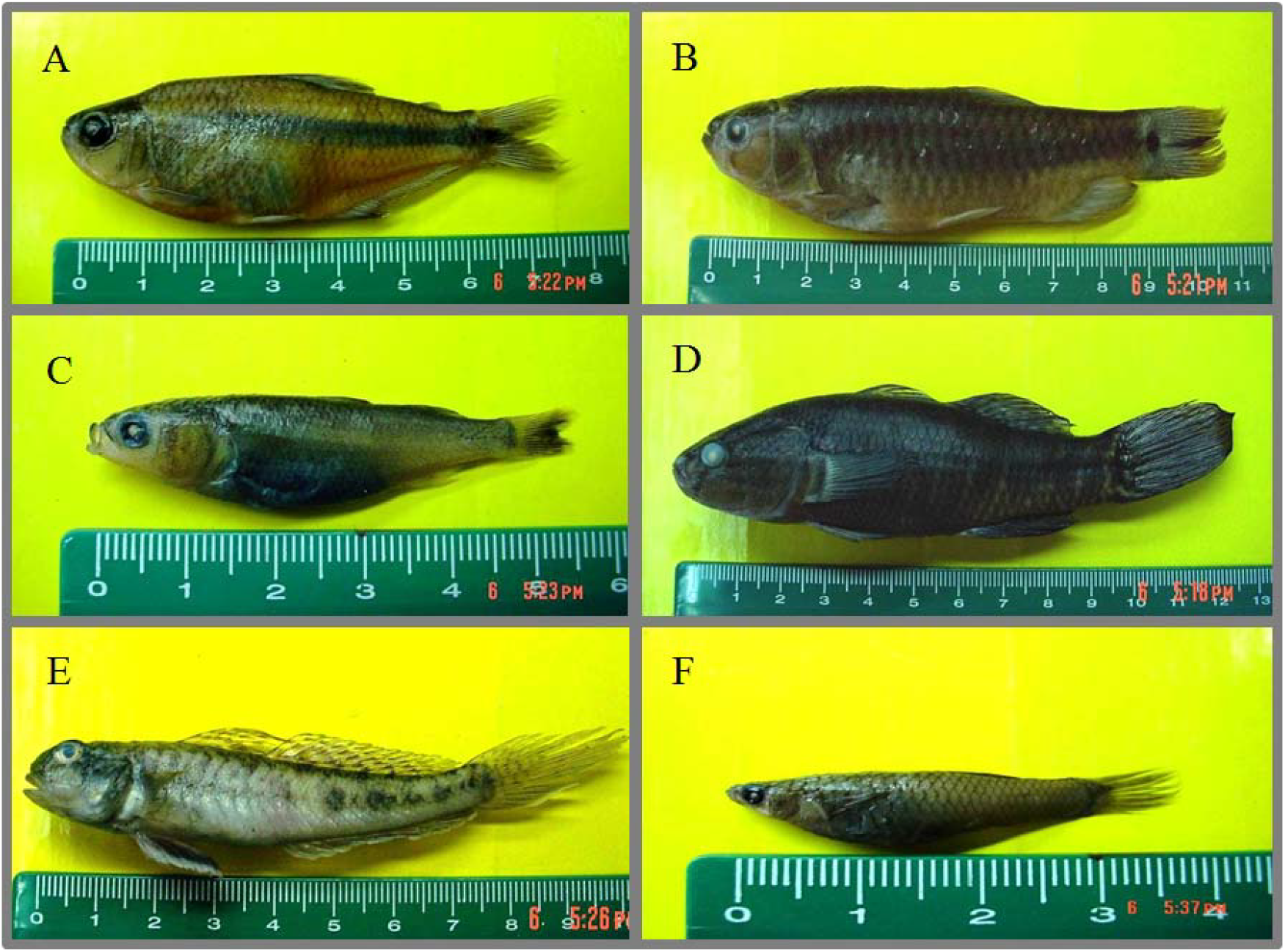
Photography of fish species collected in Totoral Lagoon, Central Peru. From A to F: *Bryconamericus peruanus, Lebiasina bimaculata, Mugil cephalus, Dormitator latifrons, Ctenogobius sagittula, Poecilia reticulate*. The measurement indicates centimeters.

In contrast, *Mugil cephalus* and the non-native but well-established *Poecilia reticulata* were among the most abundant and spatially dominant species throughout the study period. The consistent presence of *M. cephalus* near the lagoon’s outlet reflects its euryhaline nature and capacity to exploit both marine and brackish environments. Meanwhile, the success of *P. reticulata*, a species widely introduced in tropical and subtropical waters, suggests a high degree of ecological plasticity that allows it to colonize variable environments such as the Totoral Lagoon. The coexistence of native freshwater and non-native species, together with euryhaline taxa of marine origin, highlights the ecotonal character of the lagoon. This system represents a dynamic interface between continental and coastal ecosystems, acting as both a biological corridor and a temporary refuge for juvenile stages of several species. The structural complexity of its habitats, combined with tidal and seasonal fluctuations, likely drives the observed spatial segregation and abundance patterns within the ichthyofaunal community.

Strong ecological interactions, particularly trophic linkages, are a defining feature of estuarine and coastal lagoon ecosystems [19]. In the Totoral Lagoon, most species, especially *Bryconamericus peruanus*, appeared to be strongly associated with specific feeding areas. This spatial segregation may reflect distinct trophic preferences and resource partitioning within the system. Previous studies have described the predatory behavior of *B. peruanus* on the freshwater prawn *Cryphiops* (*Macrobrachium*) *caementarius* [4], suggesting that this interaction could represent a key trophic pathway in the lagoon’s food web. The abundance of *B. peruanus* in vegetated and muddy areas, where *C. caementarius* is also common, reinforces the hypothesis that these zones serve as focal points of benthic–pelagic energy transfer and trophic connectivity.

An additional noteworthy result of this study is the relatively high species richness recorded in the Totoral Lagoon despite its limited spatial extent. The number of fish species found here exceeds that reported from ecologically similar lagoons and wetlands along the central Peruvian coast (Ortega et al. 2012), even in sites of considerably larger area. For instance, studies conducted in extensive coastal wetlands of the region have documented up to 13 fish species, of which only five are native (Tovar 1977; Castro et al. 1990, 1998). The Totoral Lagoon supports an equal number of native taxa, *B. peruanus, Mugil cephalus, Lebiasina bimaculata, Aequidens rivulatus*, and *Dormitator latifrons*, in addition to the introduced *Poecilia reticulata*, the widely distributed *Cichlasoma nigrofasciatum*, and the marine-associated *Ctenogobius sagittula*. This composition underscores the lagoon’s ecotonal character, where freshwater, estuarine, and marine elements coexist, enhancing local biodiversity through species interchange and ecological overlap.

Despite its ecological importance, the Totoral Lagoon is subject to multiple anthropogenic pressures, including resource extraction, waste disposal, and habitat fragmentation. These activities have progressively altered their original configuration and threaten its ecological integrity. Nevertheless, some sectors of the lagoon remain relatively undisturbed and still harbor high species diversity and functional complexity, making them priority areas for conservation and management. The persistence of these less-impacted areas provides a unique opportunity to study the natural dynamics of lagoonal ecosystems in the central Peruvian coast. Currently, local initiatives are attempting to promote the sustainable use and restoration of the lagoon; however, its present ecological status remains poorly understood. There is limited information on the temporal variability of its physical–chemical conditions, trophic structure, and biological communities. Comprehensive, long-term monitoring programs are urgently needed to evaluate how the system responds to environmental and anthropogenic stressors.

From a broader perspective, the Totoral Lagoon exemplifies the ecological and functional significance of small coastal water bodies along the Peruvian coast. These environments act as biological corridors that connect marine and continental ecosystems, supporting the recruitment, feeding, and refuge of multiple species. Understanding their dynamics is essential for the design of integrated management strategies that reconcile biodiversity conservation with local human activities. Future research should focus on quantifying energy flows, trophic interactions, and population connectivity across these systems, thereby elucidating their contribution to the maintenance of regional fish diversity and coastal ecosystem resilience.

## Acknowledgements

I am deeply grateful to Prof. Luis Hoyos for his collaboration in obtaining the specimen photographs. I also wish to thank Prof. Roger Quiroz for providing access and support to work in Laboratory S-7, Faculty of Biological Sciences, San Marcos University.

